# Subtle perturbations of the maize methylome reveal genes and transposons silenced by DNA methylation

**DOI:** 10.1101/221580

**Authors:** Sarah N. Anderson, Greg Zynda, Jawon Song, Zhaoxue Han, Matthew Vaughn, Qing Li, Nathan M. Springer

## Abstract

DNA methylation is a chromatin modification that can provide epigenetic regulation of gene and transposon expression. Plants utilize several pathways to establish and maintain DNA methylation in specific sequence contexts. The chromomethylase (CMT) genes maintain CHG (where H = A, C or T) methylation. The RNA-directed DNA methylation (RdDM) pathway is important for CHH methylation. Transcriptome analysis was performed in a collection of *Zea mays* lines carrying mutant alleles for CMT or RdDM-associated genes. While the majority of the transcriptome was not affected, we identified sets of genes and transposon families sensitive to context-specific decreases in DNA methylation in mutant lines. Many of the genes that are up-regulated in CMT mutant lines have high levels of CHG methylation, while genes that are differentially expressed in RdDM mutants are enriched for having nearby mCHH islands, providing evidence that context-specific DNA methylation directly regulates expression of a small number of genes. The analysis of a diverse set of inbred lines revealed that many genes regulated by CMTs exhibit natural variation for DNA methylation and gene expression. Transposon families with differential expression in the mutant genotypes show few defining features, though several families up-regulated in RdDM mutants show enriched expression in endosperm, highlighting the importance for this pathway during reproduction. Taken together, our findings suggest that while the number of genes and transposon families whose expression is reproducibly affected by mild perturbations in context-specific methylation is small, there are distinct patterns for loci impacted by RdDM and CMT mutants.

## INTRODUCTION

The epigenome describes the potential for additional heritable information that can be passed on through mitosis or meiosis (Hofmeister *et al*. 2017). DNA methylation is one molecular mechanism that can provide epigenetic information. There is interest in the potential for cryptic information present in genomes that is normally silenced by epigenetic mechanisms but could be activated through epigenetic changes without requiring any genetic change.

Much is known about the mechanisms that control DNA methylation and the functional roles of DNA methylation in regulating transposon and gene expression in the model plant *Arabidopsis thaliana*. However, our knowledge of the regulating mechanisms and function of DNA methylation is much more limited in crop plants. Evidence in rice and maize suggests that major perturbations of DNA methylation disrupt development and the seeds/plants are not viable (Yamauchi *et al*. 2014; Hu *et al*. 2014; Li *et al*. 2014). Forward genetic screens for factors involved in epigenetic phenomena such as paramutation (Dorweiler *et al*. 2000; Hollick *et al*. 2005; Alleman *et al*. 2006; Hale *et al*. 2007; Jr *et al*. 2009) or transgene silencing (McGinnis *et al*. 2006) have identified several genes that are associated with DNA methylation or chromatin in maize (Hollick 2017). In addition, reverse genetic approaches have been utilized in attempts to document the function of putative methyltransferase genes or other genes associated with DNA methylation (Papa *et al*. 2001; Makarevitch *et al*. 2007; Li *et al*. 2014). To date, these mutants have provided partial reductions in DNA methylation in specific sequence contexts but no mutants with drastic reductions to genomic DNA methylation have been recovered in maize.

Surveys of natural variation for DNA methylation among diverse lines of maize have revealed many examples of differentially methylated regions (DMRs) (Eichten *et al*. 2011, 2013; Regulski *et al*. 2013; Li *et al*. 2015a). A subset of the genes located near DMRs exhibit a negative correlation between DNA methylation and gene expression (Eichten *et al*. 2013; Li *et al*. 2015a). This is primarily found at genes that have CG or CHG methylation in regions surrounding the transcriptional start site (TSS) and show qualitative (on/off) expression variation among genotypes (Li *et al*. 2015a). This suggests the potential for cryptic information in the maize genome that is epigenetically silenced in some lines but can be active due to epigenetic changes in other genotypes.

Several maize mutant lines with subtle perturbations of genomic DNA methylation have been previously identified (Li *et al*. 2014). The mutants include *mop1* and *mop3*, two mutants recovered in screens for factors required for maintenance of the paramutated state at the B’ locus (Dorweiler *et al*. 2000). The *Mop1* gene encodes an RNA-dependent RNA polymerase related to RDR2 in *Arabidopsis* (Alleman *et al*. 2006) while *Mop3* is predicted to encode the second largest subunit of RNA Pol IV (Sloan *et al*. 2014), which is allelic to *rmr6 (Erhard et al. 2009)*. Mutant alleles for the two chromomethylase genes present in the maize genome, *Zmet2* and *Zmet5* also influence context-specific DNA methylation patterns (Papa *et al*. 2001; Makarevitch *et al*. 2007; Li *et al*. 2014). These genes are likely paralogs resulting from a whole genome duplication event and are orthologous to *CMT3* from *Arabidopsis thaliana*. Previous research has found that *mop1* and *mop3* genotypes have lost CHH methylation at many genomic regions with elevated CHH, and there are changes in CG and CHG at these sites as well (Li *et al*. 2014). However, as these types of regions are quite rare in the maize genome these mutants have minimal effects on genome-wide levels of CG and CHG methylation. The *zmet2-m1* mutant and, to a lesser extent, the *zmet5-m1* mutant, result in reduction of CHG methylation. These mutants also cause reductions of CWA methylation (where W is A or T) in genomic regions with low, but detectable, CWA methylation (Li *et al*. 2014; Gouil and Baulcombe 2016). Attempts to recover double mutants for *Zmet2/Zmet5* were unsuccessful, suggesting at least partially redundant function for these paralogous genes.

Mutants for *mop1*, *mop3*, *zmet2* and *zmet5* are viable with relatively few major phenotypic changes (Dorweiler *et al*. 2000; Papa *et al*. 2001). The *mop1* and/or *mop3* mutations have been shown to play important roles in the regulation of specific maize loci (Dorweiler *et al*. 2000; Alleman *et al*. 2006; Sloan *et al*. 2014), transgenes (McGinnis *et al*. 2006) or transposable elements (Lisch *et al*. 2002; Woodhouse *et al*. 2006). Microarray profiling of gene expression has revealed evidence for altered expression of small sets of genes in studies of *mop1 (Madzima et al. 2014)* and *zmet2 (Makarevitch et al. 2007)*. There is also evidence from an RNAseq experiment for altered regulation of transposable element expression in apical meristem tissue (Jia *et al*. 2009). Transcriptome analysis of rmr6, which is allelic to *mop3*, provided evidence for a potential role in stress response (Forestan *et al*. 2016). The *rmr6* mutation appears to increase the proportion of the genome that is transcribed but has subtle effects at most loci with relatively few genes with significant changes in expression level (Forestan *et al*. 2017). However, there have not been comprehensive studies on the overlap of genes or transposons that are sensitive to mutations in different CMT or RdDM genes in maize.

Each of the mutant backgrounds used for this study has subtle effects on genomic methylation levels and can produce viable plants. There are several phenotypic abnormalities observed in *mop1* and *mop3* stocks (Dorweiler *et al*. 2000; Barber *et al*. 2012; Sloan *et al*. 2014) although the penetrance in multiple backgrounds has not been well characterized. We sought to determine if the subtle changes in DNA methylation in these mutants would reveal genes or transposons that are sensitive to these shifts in DNA methylation or chromatin. A limited number of genes and transposon families exhibit altered expression in these genotypes. A subset of these genes have high levels of DNA methylation in wild-type that are reduced in the mutant genotypes. Many of these genes exhibit natural variation for DNA methylation and gene expression. This provides evidence that the natural variation at these genes is due to epigenetic rather than genetic variation and highlights cryptic information present in the maize genome that could be accessible through alterations to the epigenome.

## MATERIALS AND METHODS

### Biological materials

All the mutant and wild type samples used in this study are listed in Table S1 and SRA accession numbers are listed for each dataset. Tissue for RNA and DNA isolations was collected from three biological replicates. Plants were grown in standard greenhouse conditions for 20 days to reach the V3 stage. The 2^nd^ and 3^rd^ leaves were collected individually for each seedling. The 2^nd^ leaf was used to isolate DNA for genotyping, and the 3^rd^ leaf was used for RNA isolation and sequencing. For each biological replicate, 4-6 seedlings were pooled.

### Library preparation and sequencing

Total RNA was isolated using the TRIZol reagent following the manufacturer’s protocol. RNA was quantified using RiboGreen and 3 μg total RNA was used to construct libraries using TruSeq strand-specific kit (Illumina) following manufacturer’s suggestions. The final library was quantified using PicoGreen and twelve libraries were pooled per Illumina lane. Library quality was checked using Agilent Bioanalyzer. Sequencing was performed on HiSeq2500 using 2 x 50 bp mode.

### Gene expression analysis

Trim_glore was used to trim low-quality base from the 3’ end of the reads, as well as to remove adapters. Reads that passed quality control were mapped to B73 version 4 genome (Jiao *et al*. 2017) using Tophat2 (Kim *et al*. 2013), allowing at most 1 mismatch (-N 1) and the expected inner distance between mate pairs of 200 bp (-r 200). Reads that are properly paired and uniquely mapped were filtered out using samtools (-f 0x0002 −q 50). HTSeq (Anders *et al*. 2015) was used to summarize the number of reads mapped to each V4 gene model with the union mode, generating a matrix of count values for each gene in each genotype.

Raw read counts were input into DEseq2 (Love *et al*. 2014) to perform differential expression analysis. Pair-wise comparisons were made between each mutant and the appropriate wild type. Genes with a FDR value of < 0.05 and log_2_(FoldChange) > 1 were called differentially expressed genes. Detailed analysis was restricted to genes with consistent DE calls in at least two mutant contrasts in the same pathway (RdDM or CMT). Genes were considered expressed if at least 3 replicates in the libraries described had an RPM (reads per million) value > 1.

### TE expression analysis

B73v4 (Jiao *et al*. 2017) TE annotation was modified to remove helitrons and the file was resolved using RTrackLayer in R so that each base of the genome was assigned to only a single TE. Exon regions were masked from the TE file using Bedtools (Quinlan and Hall 2010) subtract. Gene annotations were added to this modified TE annotation file, and mapped reads were assigned to features using HTSeq (Anders *et al*. 2015). A custom script was used to read through the HTSeq sam output, assigning unique-mapping reads to individual TE elements and multi-mapped reads to TE families if mapped positions hit only a single TE family. Unique and multi-mapped reads were combined for per-family expression counts, and RPM values were calculated by normalizing to the number of gene reads plus TE family reads in each library. All reads mapped to gene annotations plus TE annotations were excluded from the TE expression analysis. Differentially expressed TE families were determined using DESeq2 (Love *et al*. 2014) using a log_2_(FoldChange) cutoff of 1 and FDR adjusted p-value cutoff of 0.05. Detailed analysis was restricted to TE families with consistent DE calls in at least two mutant contrasts in the same pathway (RdDM or CMT).

### WGBS data analysis

The WGBS datasets used in this study are detailed in Table S2 and SRA accession numbers for each sample are provided. One μg DNA was used to prepare libraries for whole-genome bisulfite sequencing using the KAPA library preparation kit. DNA was sheared to a peak between 200-250 bp. End repair was performed to make blunt-ended fragments, followed by adding base A to the 3’ end, and adapter ligation. Size selection was performed to enrich library with a size between 250-450 bp. Bisulfite conversion was then carried out using Zymo DNA methylation lightning kit according to user’s manual. Finally, library was enriched using PCR amplification. Library quality was checked using the Agilent Bioanalyzer. Library quantification was performed with qPCR before sequencing. Sequencing was performed on HiSeq2000 with paired end 100 cycles.

Analysis was performed as previously described (Li *et al*. 2015a; Song *et al*. 2016). Read quality was checked with FASTQC, adapters and low-quality bases at the 3’ end of each read were trimmed using Trim_glore. The high quality reads were mapped to B73 V4 genome (Jiao *et al*. 2017) using BSMAP (Xi and Li 2009) allowing at most 5 mismatches. Only properly paired reads with unique mappings were kept and used for calling DNA methylation. Methylation calls were performed using the methratio.py script from BSMAP. Finally, DNA methylation in each context (CG, CHG, CHH) was summarized for each 100-bp non-overlapping tile of the 10 maize chromosomes.

### DMR calling

DMRs were called using previously described criteria (Li *et al*. 2015a). Briefly, each 100-bp tile with > 6 CG/CHG sites, > 2X coverage and > 60% difference for CG/CHG were compared. CHH DMRs were called using the same coverage and site number criteria, but with a requirement for <5% CHH in one genotype and >25% CHH in another genotype, reflecting the low level of CHH methylation in the maize genome.

### mCHH Islands

High CHH bins were called genome-wide by requiring CHH methylation over 25% with at least 10 informative counts per bin. Genes and TEs were considered associated with a mCHH island if when at least one high methylation bin was identified within the gene or in the 2 kb region surrounding the gene.

### Data availability

All data used in this study are deposited at the NCBI Sequence Read Archive (SRA). Accession numbers for all libraries are listed in Table S1.

## RESULTS

### Alteration of gene and transposon expression in maize mutants with perturbed methylomes

RNAseq was used to perform transcriptome profiling for several maize lines carrying mutations in genes encoding CMT (this study) or RdDM components (Gent *et al*. 2014; Li *et al*. 2015b). Together these factors are expected to be responsible for the majority of CHG and CHH methylation in the maize genome. For CMT genes, three biological replicates of seedling leaf tissue were profiled for mutations in two different genes, with multiple alleles utilized for one of the genes (Table S1). In addition, for the RdDM genes we analyzed seedling leaf tissue for *mop1* and *mop3* (Li *et al*. 2015b), along with immature ear tissue for *mop1* (Gent *et al*. 2014). The genetic background, read number and accession information for each sample is provided in Table S1.

The expression of individual genes was estimated from the RNAseq data for each sample. Differentially expressed (DE) genes in each mutant line (relative to the appropriate control) were identified using DESeq2 followed by a requirement for a minimum of 2 fold-change and an FDR value of less than 0.05 (Table S2). The observed differences in gene expression in the mutant lines could be direct effects of the mutation on expression, indirect effects caused by direct targets, or could be the result of introgressions of linked loci that contain cis-regulatory variation. The number of genes in each 2 Mb bin with differential expression was assessed throughout the genome (Figure S1). For mutations that were identified in one background and then backcrossed into another background (*zmet2-m1, zmet2-m2, zmet5, mop1*), there were often a cluster of DE genes surrounding the locus of the mutation itself. These regions often included similar numbers of up- and down-regulated genes. For the other mutation (*mop3*) that was not backcrossed into another genetic background, there is less evidence for expression changes at linked genes (Figure S1). Based on these results we omitted DE genes located within 40 Mb of the mutation in subsequent analyses.

A principle component analysis was performed using all DE genes to cluster samples used in this study (Figure 1). This reveals that the major sources of differences are tissue and genetic background, as demonstrated, respectively, by the relatively larger differences for *mop1-ear* and *mop3* which was not introgressed into B73 (Figure 1A). When comparing only leaf libraries in the B73 background, few genotypes were substantially different from the wild-type controls, suggesting limited changes to transcript levels induced by each mutation (Figure 1B). The number of differentially expressed genes in each mutant genotype was highly variable (Figure 1C). In most cases the homozygous mutant individuals exhibit more up-regulated genes than down-regulated genes, which is compatible with the concept that the CMT and RdDM genes normally provide silencing activities.

**Figure 1:**
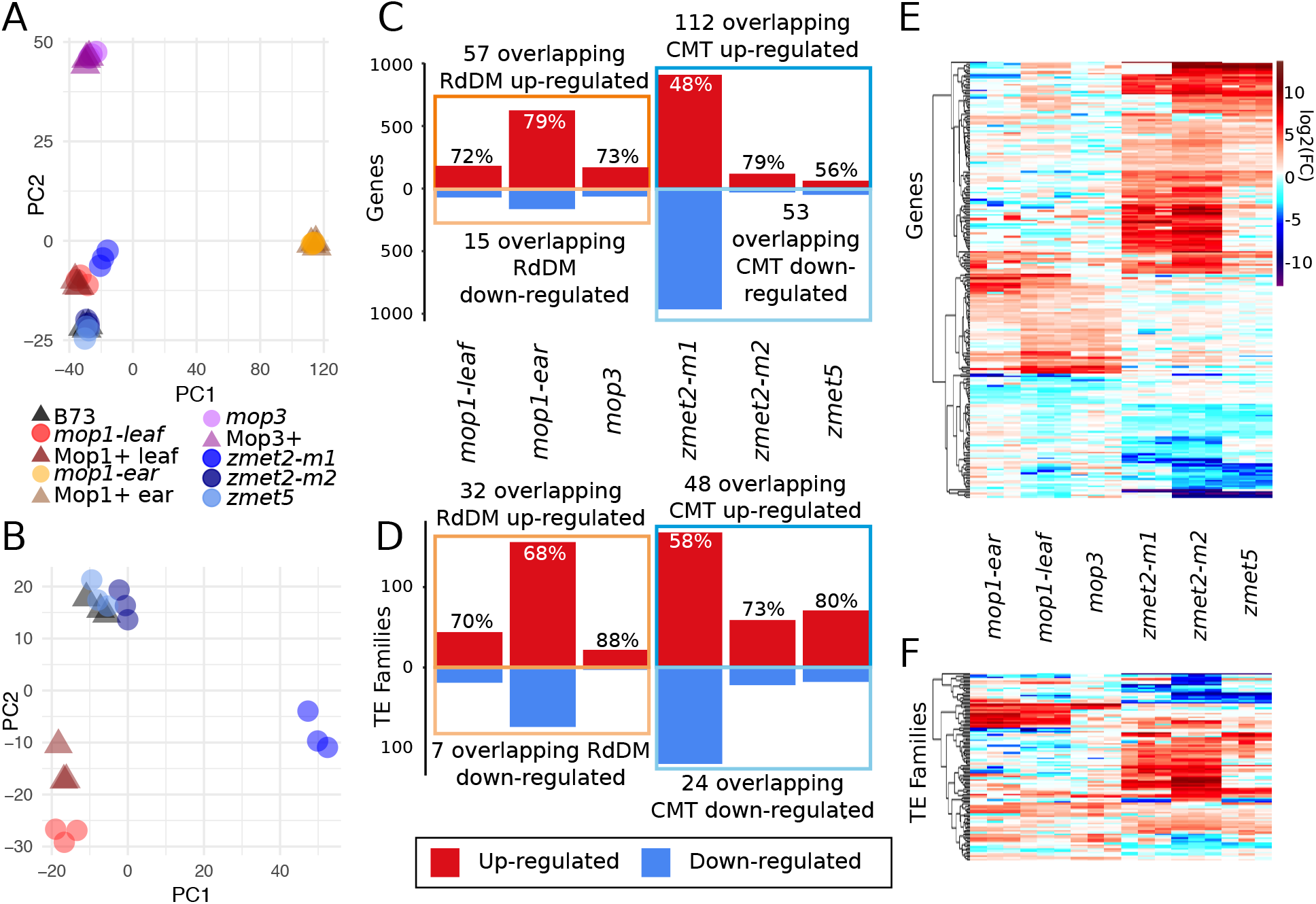
Summary of differentially expressed genes in mild methylation mutants. A-B: A principal component analysis (PCA) was performed using log_2_(RPM+1) expression values for genes that are DE in at least one mutant line relative to the appropriate control. (A) The full set of samples used for this study were assessed and we found that samples in other genetic backgrounds (*mop3* and *Mop3*) or tissues (*mop1* ear and *Mop1* ear) have the highest level of variation. (B) A second PCA was performed using only samples in the B73 genetic background assessed in leaf tissue. WT samples are denoted with triangles and mutants with circles. C-D: The number of up- (red) and down-regulated (blue) genes (C) and TEs (D) is shown for each mutant relative to the appropriate wild-type control. The percent of DE genes or TE families that are up-regulated is marked above each bar, and the number of genes or TEs with consistent changes in two or more mutants from the same pathway are labeled. E-F: Each of the genes (E) or TE families (F) that are DE in at least two CMT or RdDM mutants were used to perform hierarchical clustering using the Euclidean method and the log_2_ of the fold-change relative to wild-type is visualized with a heat map.

Transposable elements (TEs) comprise a large portion of the maize genome, and typically have high levels of CG and CHG methylation, with CHH methylation peaks at the edges of some TE families. There are two classes of TEs, Class I (retrotransposons) and Class II (DNA transposons), which transpose either through a copy-and-paste mechanism requiring an RNA intermediate (retrotransposons) or through a cut-and-paste mechanism (DNA transposons) (Wicker *et al*. 2007). Within each class are several orders divided into superfamilies, distinguished by structural and protein-coding features. Families within each superfamily are defined by sequence identity, and each family can contain any number of individual TE elements (Jiao *et al*. 2017). Individual TE elements are defined at a single location within a genome and are associated with a family, superfamily and class. We sought to document how minor perturbations to the methylome impacted expression of TEs. Due to the highly repetitive nature of TE sequences, we assessed per-family levels of expression by mapping RNA-seq reads to the genome, reporting up to 20 best hits for each read using Tophat2. Per-family read counts were determined by summing unique mapping reads (to a specific element) and multi-mapping reads that align to only a single TE family. Overall, the total portion of RNAseq reads that map to TE families is not significantly higher in the mutants than in wild-type plants suggesting a lack of genome-wide activation of TEs in these mutants (Figure S2). In order to assess expression of individual TE families in each genotype, per-family expression was normalized by dividing the family counts by the total number of reads in the library assigned to either TE families or genes, generating an RPM estimate. Using this approach we were able to detect expression of 1,694 TE families in at least one of the genotypes used for this study. A relatively small number of DE TE families (log_2_FC > 1, FDR < 0.05) were identified in each mutant (Table S3; Figure 1D). Consistent with the role of DNA methylation in silencing TEs, more families were identified as up-regulated rather than down-regulated in mutants compared with WT controls. However, the majority of the TE families expressed in these libraries do not exhibit significant changes in expression level in CMT or RdDM mutants in maize.

There is a significant overlap in the number of genes and TEs that exhibit consistent changes in gene expression in at least two samples of CMT mutants (*zmet2-m1/zmet2-m2/zmet5*) or RdDM mutants (*mop1/mop3*) (Figure S3). In order to understand the reproducible effects of these pathways on expression, we focused our analyses on the set of 237 genes and 104 TE families that exhibit consistent up- or down-regulation in multiple CMT or RdDM mutants. Hierarchical clustering of the expression level of these genes or TE families in all samples reveals evidence for many consistent changes in expression within a pathway (Figure 1E-F). Within CMT mutants, many genes show shared expression changes between *zmet2-m1* and *zmet2-m2*, though there is a smaller subset of genes primarily shared between *zmet2-m2* and *zmet5* (Figure 1E). Although both *zmet2-m1* and *zmet2-m2* are predicted to encode loss of function alleles, there are some genetic differences in the behavior of these alleles (Papa *et al*. 2001; Makarevitch *et al*. 2007; Li *et al*. 2014). The *zmet2-m1* mutation exhibits partial dominance that may reflect dominant negative action of the protein that could be produced from this allele (Papa *et al*. 2001). Plants that are homozygous for *zmet2-m1* have the greatest loss of CHG methylation and this could result from influence of the *ZMET2* protein product on functional *ZMET5* protein. The *mop1* and *mop3* seedling leaf samples have a number of examples of consistent up-regulation but fewer examples of consistent down-regulation, consistent with the greater number of up-regulated than down-regulated genes in RdDM mutants in general.

### Some genes that are up-regulated in CMT mutants exhibit high CHG methylation levels

The differential expression observed in each mutant background could result from direct changes in DNA methylation or chromatin at these loci or could result from indirect effects due to secondary effects from genes that are direct targets. Given that many of these mutants are expected to affect DNA methylation levels, we might expect that wild-type plants would contain high levels of DNA methylation for genes that exhibit increased expression in the mutants. The context-specific DNA methylation profiles were assessed in wild-type B73 for genes that were up- or down-regulated compared with all expressed genes (Figure 2A). Genes that are differentially expressed in RdDM mutants exhibit slightly higher levels of CHH methylation upstream of the transcription start sites relative to all expressed genes. The genes that are down-regulated in CMT mutants do not show unusual patterns of DNA methylation. In contrast, genes that are up-regulated in the CMT mutants exhibit distinct patterns of CG and CHG methylation within gene bodies relative to other expressed genes (Figure 2A). Among the 112 genes up-regulated in CMT mutants, approximately half have high (>50%) and half have low (<20%) methylation in the CG and CHG contexts (Figure 2B-C). In contrast, only ~4% of all expressed genes have high CHG methylation in the same region. While a small number of genes with high methylation overlap annotated TEs, most of the genes in this subset do not, suggesting that this genic methylation is not solely due to nearby TEs. In wild-type samples, genes with high CHG methylation in the gene body also have high CG methylation (Figure 2D). However in *zmet2-m1* mutants, CHG methylation for these genes is reduced, with few examples of a corresponding reduction in CG methylation (Figure 2D-E). As in the examples *(Zm00001d045627* and *Zm00001d021982*) shown in Figure 2F, these genes have high levels of CG and CHG methylation in wild-type B73, and the reduction in CHG methylation in *zmet2-m1* mutants is associated with increased expression, suggesting that CMT-dependent silencing of these genes depends on CHG but not CG methylation.

**Figure 2:**
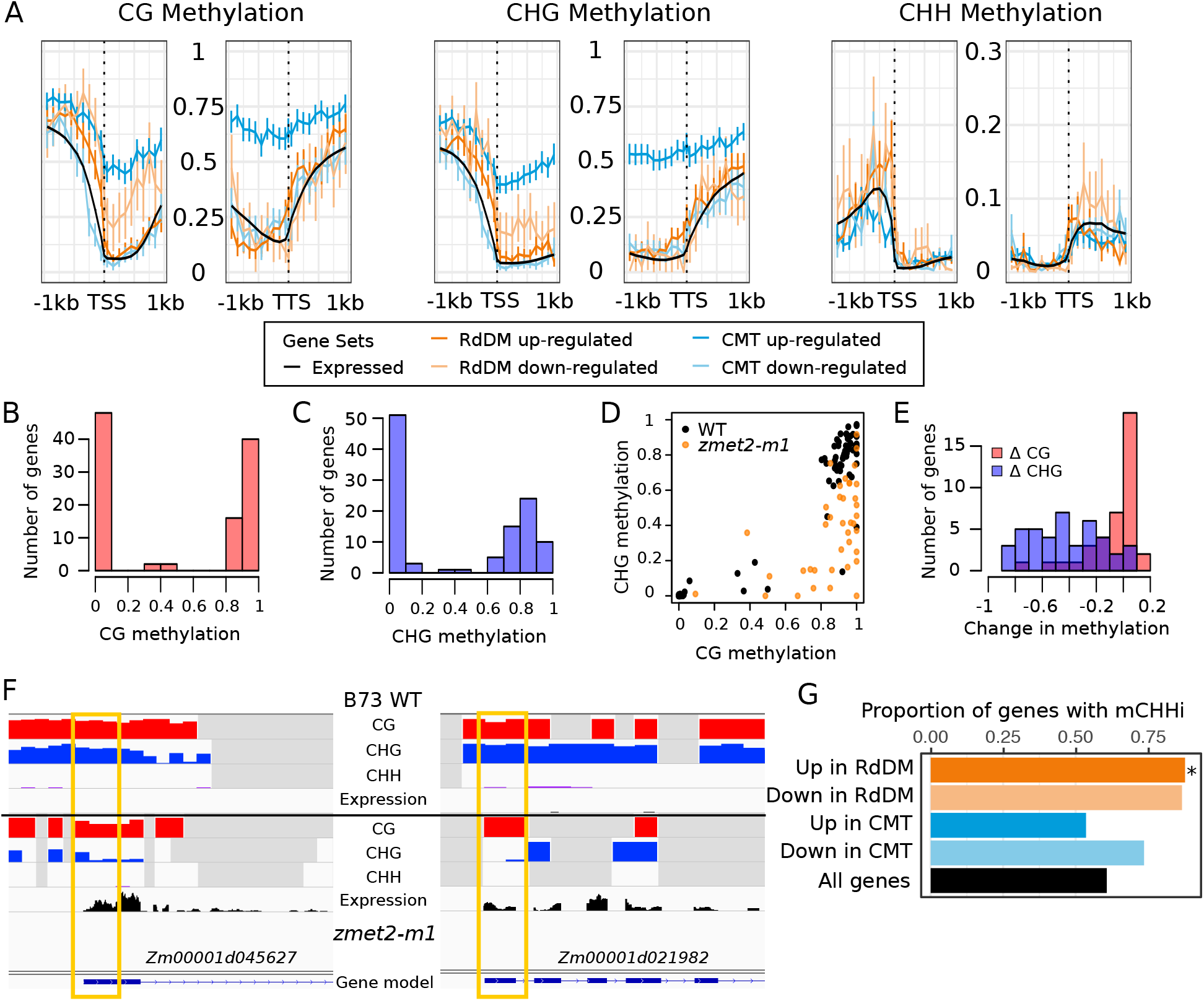
Methylation profiles of DE genes. A. The metaprofile of DNA methylation levels in wild-type B73 seedling leaf tissue was assessed for different sets of DE genes. The DNA methylation levels surrounding the TSS and TTS were plotted for all expressed genes and genes DE in CMT (blue) and RdDM (orange) mutants. The three panels show the levels of CG, CHG, and CHH methylation, with the y-axis showing DNA methylation levels. Error bars represent standard error. B-C Histogram of the number of genes with different methylation levels in the first 400 bp downstream of the TSS in the CG (B) and CHG (C) contexts, showing a bimodal distribution of methylation values. D. Wild-type methylation levels in the CG and CHG contexts are correlated. *zmet2-m1* mutant methylation data is shown for those genes with high (>50%) CHG methylation in wild-type (orange dots), showing a loss of CHG but not CG methylation in the mutant. E. Histogram of the difference between mutant and WT methylation in the CG (red) and CHG (blue) contexts for those genes up-regulated in CMT mutants that have WT CHG methylation >50%. F. IGV view of two up-regulated genes in CMT mutants: *Zm00001d045627* and *Zm00001d021982*, which have high CG and CHG methylation in WT and reduced CHG methylation near the TSS (yellow box) in *zmet2-m1* mutants. G. The proportion of genes within 2 kb of mCHH islands (mCHHi) is shown for different sets of genes. The black bar shows the proportion of all genes with a mCHH island while the other bars show the proportion of genes with altered expression in specific mutant backgrounds that have mCHH islands, and * denotes significantly higher proportion than expected relative to all genes (p-val < 0.01, chi squared test).

### Genes that are up-regulated in RdDM mutants are enriched for being near mCHH islands

A large number of maize genes (~60%) have been associated with the presence of a region of elevated CHH in the promoter region, termed a mCHH island (Gent *et al*. 2013; Li *et al*. 2015b). Genes with mCHH islands are enriched for high expression and the mCHH island often occurs at the edge of the TE nearest these genes (Li *et al*. 2015b). These mCHH islands may form important boundaries that could protect TE heterochromatin from the influence of genes (Li *et al*. 2015b) and may also be important for long-distance interactions (Rowley *et al*. 2017). The methylation within these mCHH islands requires *mop1* and *mop3* (Li *et al. 2014, 2015b)*. We sought to determine if the genes that exhibit altered expression in *mop1* and *mop3* are enriched for the presence of mCHH islands. Genes that are up- or down-regulated in the RdDM mutants are enriched for the presence of mCHH islands, but this is only significant for the up-regulated genes with 87.8% having a mCHH island within 2kb of the gene, compared to 64.6% of all expressed genes (Figure 2G). The fact that both RdDM up- and down-regulated genes are often near mCHH islands could be due to the fact that the mCHH island may provide long-range interactions (Rowley *et al*. 2017) that could have either positive or negative influences on gene expression.

In some cases, the mCHH island itself may result in transcriptional regulation. Work in *Arabidopsis* has noted a positive feedback loop involving DNA methylation levels and expression of the demethylase enzyme *ROS1* such that reduced levels of DNA methylation result in lower *Ros1* expression but increased methylation is associated with elevated *Ros1* expression (Williams *et al*. 2015). Reduced expression of maize DNA glycosylases has also been observed in several transcriptome datasets of maize RdDM mutants (Williams *et al*. 2015; Erhard *et al*. 2015). We find that one maize gene with sequence homology to *Ros1*, *Zm00001d038302*, showed significantly reduced expression in the *mop1* and *mop3* mutants and has a strong mCHH island in several inbred lines (Figure S4). This provides evidence to support a requirement for RdDM and CHH methylation in the proper control of this gene in maize.

### Genes regulated by CMT are enriched for natural DMRs

The genes that are sensitive to mutations in CMT or RdDM components may reflect examples of natural variation for epigenetic regulation. Indeed, an earlier study has found that many of the genes influenced by *Zmet2* exhibit variable expression patterns in different maize inbreds (Makarevitch *et al*. 2007). We used WGBS data from B73 and 17 other diverse maize inbreds to document natural variation for DNA methylation among maize inbreds. Differentially Methylated Regions (DMRs) were identified in all three contexts (CG, CHG, and CHH) between B73 and the other inbreds. More than 200,000 DMRs were called in the CG and CHG contexts, with over 50,000 DMRs in the CHH context. Each maize gene was classified based on whether there was a DMR within 200 bp of the transcription start site for each of the three sequence contexts. A relatively small portion (~3-7%) of maize genes have CG, CHG or CHH DMRs near the promoter (Table S2). We proceeded to assess whether naturally variable DMRs were more prevalent near genes that exhibit altered expression in CMT or RdDM mutants (Figure 3A). Genes that are up- or down-regulated in CMT mutants exhibit a significant enrichment for CG and CHG DMRs in their promoter regions. RdDM up-regulated genes are significantly enriched for having CHH and CHG DMRs and also show an enrichment (though not significant) for CG DMRs near the promoter (Figure 3A).

**Figure 3.**
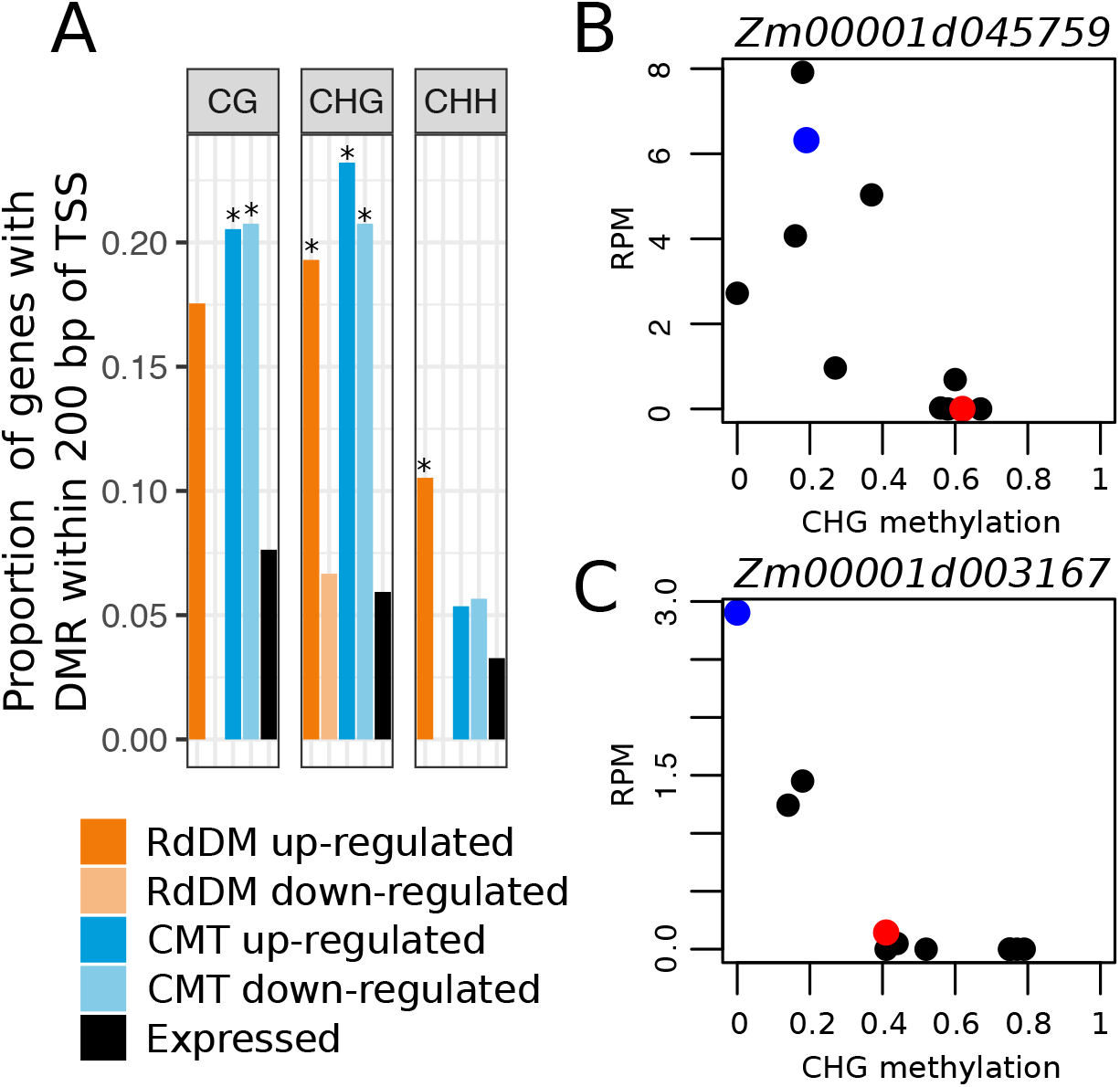
Natural variation for methylation level. A. DMRs in diverse genotypes located within 200 bp the TSS for all genes and for genes with differential expression in either CMT or RdDM mutants. Significant enrichment compared with expressed set is denoted with * (p-value < 0.01, chi squared test). B-C.Examples of genes up-regulated in CMT that have negatively correlated expression and CHG methylation at the bin overlapping the TSS. Data points show values for B73 (red), *zmet2-m1* mutants (blue), and 9 diverse genotypes (B97, CML322, HP301, IL14, Mo17, Oh43, P39, Tx303, and W22, black).

This suggests that many of the genes with altered expression in CMT or RdDM mutations may have pre-existing natural variation for DNA methylation that would affect expression levels in maize populations. RNAseq data from leaf tissue for ten of the inbred lines with WGBS data was utilized to determine whether there was a significant association (p.value < 0.05, pearson correlation) between context-specific methylation level at the DMR and gene expression levels. We found that nearly 50% of the genes that exhibit altered expression in CMT or RdDM mutants that are located near DMRs had natural variation for gene expression levels that was significantly associated with DNA methylation levels. Two examples of CMT up-regulated genes that exhibit significantly correlated expression and CHG methylation at the bin overlapping the TSS among diverse lines are shown in Figure 3B-C. In wild-type maize inbred lines we see two classes with respect to expression level and CHG methylation levels at the DMR near the TSS. In one group of lines, including B73, the DMR is highly methylated and the gene is transcriptionally silent. In the other group of genotypes (and in B73 *zmet2-m1* mutant lines - blue dots) the DMR has low methylation and the gene is expressed. Although we were interested in performing a similar analysis for natural variation in TE methylation and expression we were not able to assess this due to the highly polymorphic nature of TEs among different maize lines and the lack of de novo assemblies for other genotypes.

### Properties of TEs with altered expression

There are 104 TE families with altered expression in mutants that perturb RdDM or CMT components in maize. These included 32 families up- and 7 families down-regulated in at least two contrasts of RdDM mutants, and 48 families up- and 24 families down-regulated in at least two contrasts of CMT mutants (Figure 1D). There are examples of both class I (specifically Long Terminal Repeat or LTR) and class II (specifically Terminal Inverted Repeat or TIR) TE families that exhibit altered expression in both RdDM and CMT mutants (Figure 4A). Most families with varied expression were small (< 10 members), consistent with the genome-wide distribution (Figure 4B). The relative age of the LTR families that have altered expression was assessed to determine if they were particularly young or old families (Figure 4C). The relative ages of LTR transposons can be approximated by comparing the sequence similarity of the two LTR sequences. Since LTR sequences must be identical upon initial integration, a higher LTR similarity denotes younger TE insertions. LTR families that are up-regulated in RdDM mutants are enriched for younger LTR elements when compared with the distribution of ages present genome-wide. We also tested the mean GC content of TEs within families to test whether families depleted in cytosines are more susceptible to subtle perturbations in methylation, as is the case for the *ONSEN* family in *Arabidopsis* (Cavrak *et al*. 2014). TE families up-regulated in RdDM mutants do have a slightly lower GC content on average, though it is not clear if this change alone is sufficient to cause the expression changes (Figure 4D).

**Figure 4.**
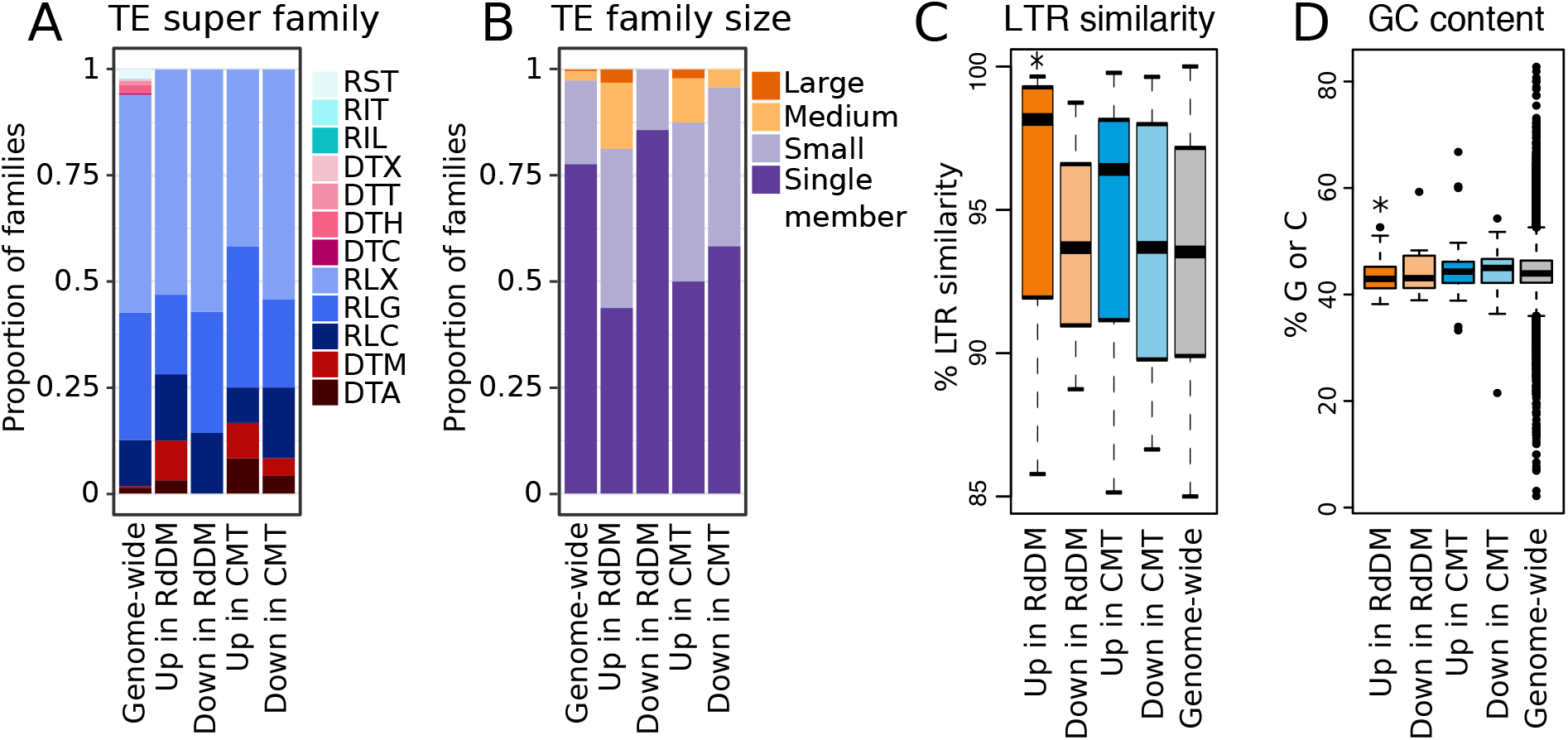
Attributes of TE families with altered expression in methylation mutants. A. TE super family membership for TE families genome-wide and with varied expression in mutants, where RST = SINE, RIT = LINE, RIL = LINE-L1-like, DTX =TIR-unclassified, DTT = TIR-Tc1/Mariner, DTH = TIR-PIF/Harbinger, DTC = CACTA, RLX = LTR-unclassified, RLG = LTR-Gypsy, RLC = LTR-Copia, DTM = TIR-Mu, and DTA = TIR-hAT. B. Size distribution for TE families genome-wide and with varied expression in mutants, where small: 2-9 members, medium: 10-99 members, and large: >= 100 members. C. Boxplot of the average LTR similarity per-family for LTR TE families genome-wide, along with those LTR families with differential expression in methylation mutants. D. Boxplot of the average GC content per-family for TE families genome-wide and with expression changes in the mutant. * denotes significant deviation from the mean for all TE families (p-value < 0.01, t-test).

We sought to further document the properties of these TE families through analysis of their expression in nearly 100 developmental tissues or stages of B73. During typical development, approximately 3,400 TE families are expressed in at least one tissue or stage. There are 5 TE families up-regulated in RdDM mutants and 18 TE families up-regulated in CMT mutants that are not expressed in any tissue or developmental stage assessed. The other families of TEs that are up-regulated in RdDM mutants (25 families) or CMT mutants (27 families) were assessed to determine if they exhibit distinct patterns of expression. Interestingly, approximately one third of the TE families that are up-regulated in RdDM mutants show higher expression in the endosperm than other tissues (Figure 5A). In contrast, the TE families up-regulated in CMT mutants do not show any evidence for higher expression in a particular tissue type. The enrichment for endosperm expression in TE families up-regulated in RdDM mutants does not extend to genes up-regulated in the mutants and cannot be simply attributed to lower expression of the *Mop1* and *Mop3* genes in these tissues (Figure 5). This result highlights the potential for some TEs to escape RdDM-based silencing in endosperm, where dynamic changes to DNA methylation may reinforce TE silencing in the embryo (Martínez and Slotkin 2012; Gehring 2013; Wang *et al*. 2015; Dong *et al*. 2017). Meanwhile, both genes and TE families susceptible to mis-regulation in CMT mutants are less often expressed across development, consistent with the greater developmental stability of CHG methylation over CHH methylation (Kawakatsu *et al*. 2016, 2017; Narsai *et al*. 2017; Bouyer *et al*. 2017).

**Figure 5.**
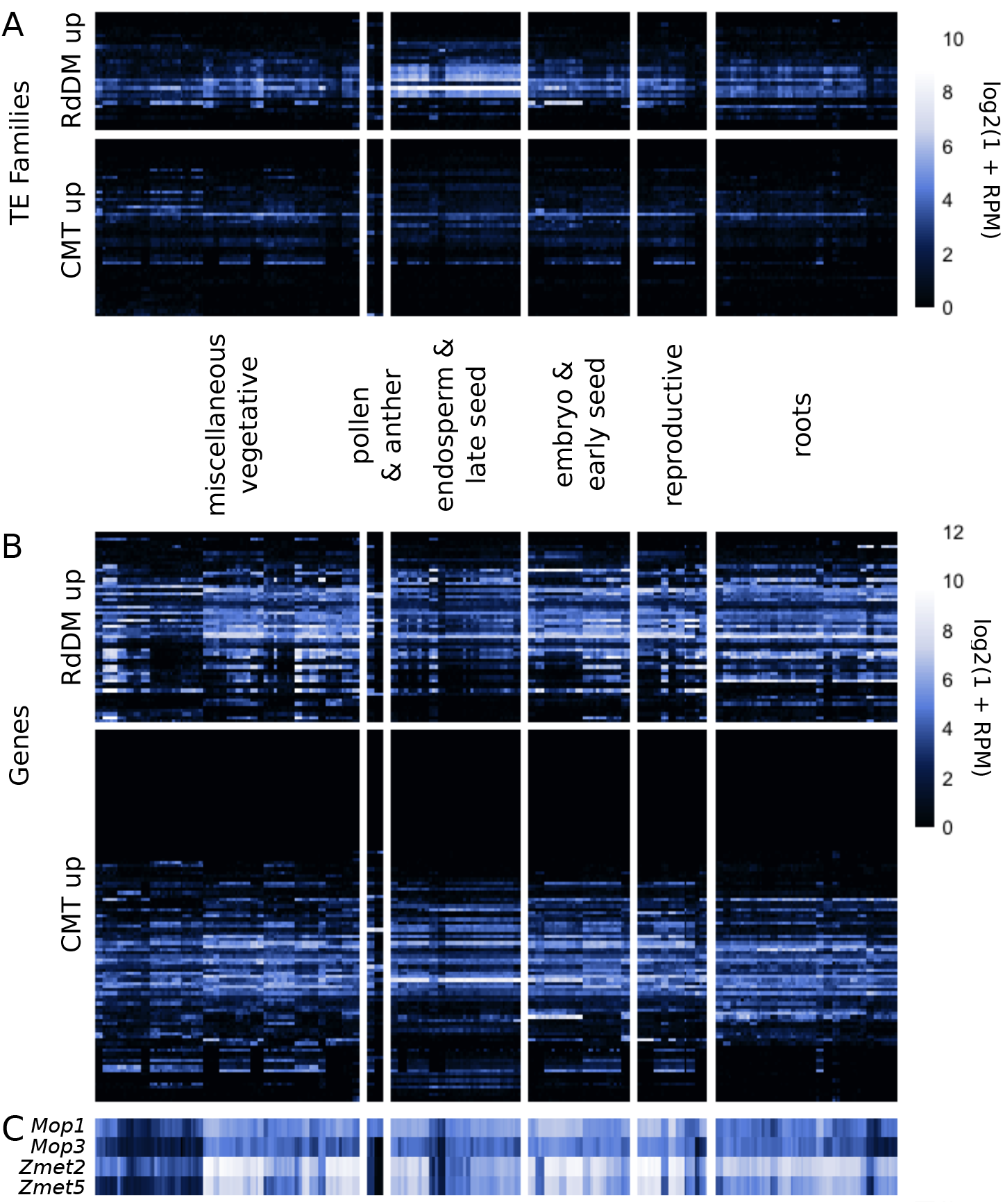
Developmental expression of TE families (A) and genes (B) up-regulated in RdDM and CMT mutants, along with the typical expression of genes mutated in this study (C), where rows show TE families or genes and columns show RNA-seq libraries. Developmental samples are grouped by tissue type, with seed samples split into two clusters based on relative contribution of endosperm: endosperm & late seed (12+ days after pollination) and embryo & early seed (up to 10 days after pollination). For full list of tissue assignments and RNA-seq library accession numbers, see Table S1. Approximately one third of TE families up-regulated in RdDM mutants have higher expression in endosperm than other tissues across development, a pattern not observed for genes. In contrast, many more TE families and genes up-regulated in CMT mutants are never or lowly expressed during typical development.

### Evidence for locus-specific and coordinated changes in expression of TEs

While per-family analysis is useful in capturing additional expression dynamics of repetitive transposable elements, the expression of individual elements can be influenced by a variety of location-specific attributes such as methylation levels and proximity to genes as well as family-level attributes such as binding motifs and nucleotide content. We were interested in documenting the relative behavior of different elements within the same family to understand whether the changes in expression of TEs were occurring in an element-specific or family-wide manner. Coordinate changes in expression could indicate the importance of RdDM or CMT for family-wide regulation while element-specific changes could reflect influences at particular loci. A set of TE families with <10 elements that had altered expression and for which at least 50% of the reads could be uniquely assigned to specific element were identified and used for analysis of coordinate versus locus-specific expression (Tables S3, S4). The unique mapping reads for these families were used to evaluate element-specific expression. Half of the testable TE families had expression of a single member of the family indicating locus-specific changes (examples in Figure 6A, C). In the other half of the TE families there was evidence for expression changes for multiple elements of the same family (Figure 6B, D). This suggests at least some level of coordinate regulation of multiple members of the family by CMT or RdDM pathways. However, even in examples of coordinate expression a single element accounted for the vast majority of unique reads mapping to the family. Examples of both locus-specific and coordinate changes in expression for both CMT and RdDM mutants were found but we were not able to assess enough families to determine if there was any enrichment for the type of regulation for these two silencing pathways.

**Figure 6:**
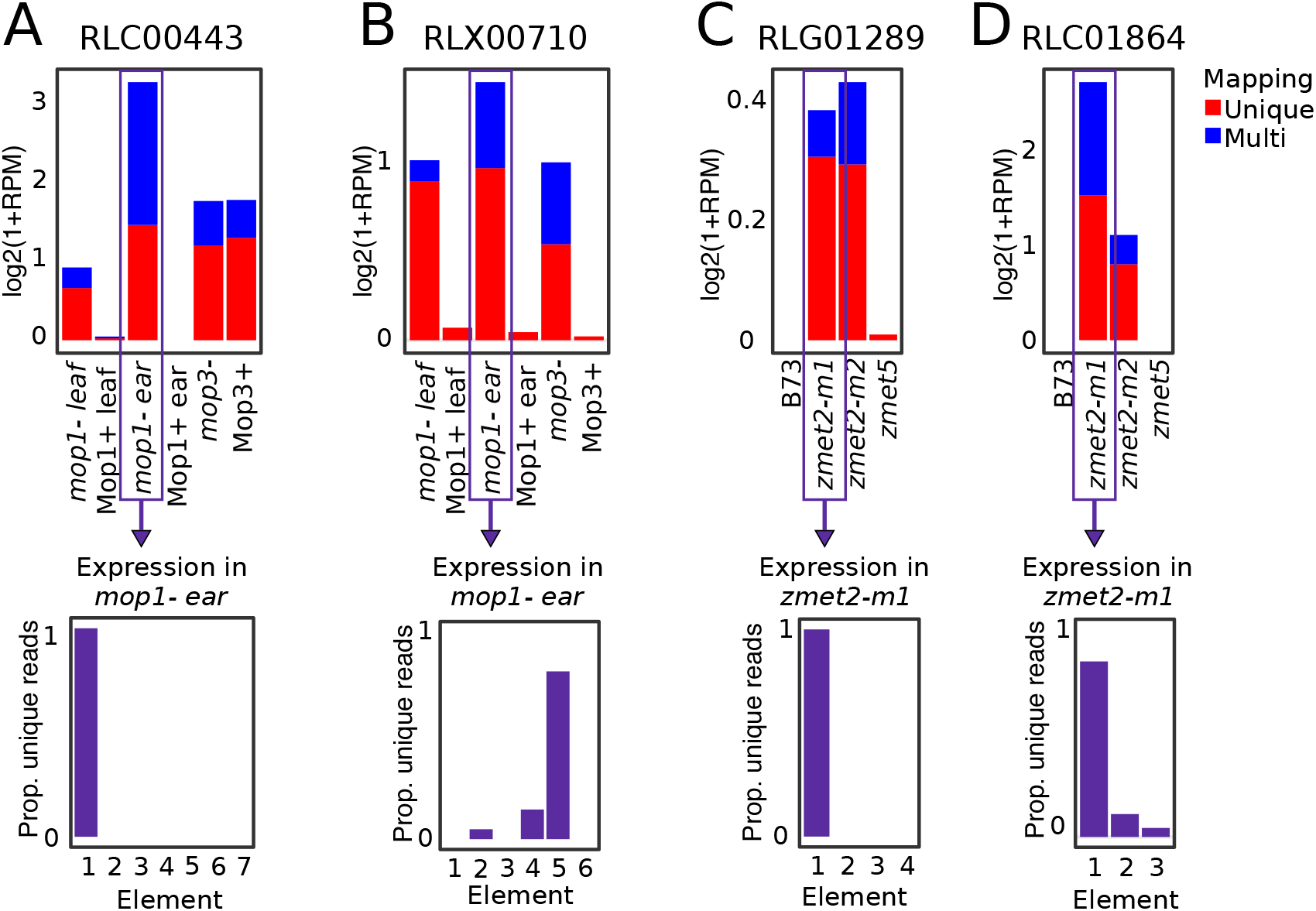
TE families up-regulated in methylation mutants can have expression of a single element or multiple elements. A-B Show two LTR families up-regulated in RdDM mutants, and C-D show two LTR families up-regulated in CMT mutants. All families have both unique (red) and multi-mapped (blue) reads. The proportion of unique-mapping reads assigned to each element is shown for a representative library. Unique-mapping reads showed confident expression of only a single member of a family (A and C) or coordinated expression of more than one member of a family (B and D).

## DISCUSSION

Maize has been a model system for the discovery of several epigenetic phenomena such as imprinting (Kermicle 1970; Kermicle and Alleman 1990), paramutation (Brink 1956; Chandler 2007; Hollick 2017) and transposon silencing (Chandler and Walbot 1986; Chomet *et al*. 1987). An unresolved question is whether epigenetic regulation plays important roles in quantitative trait variation beyond handful of well characterized loci. Our ability to document the full role for epigenetic regulation and DNA methylation has been limited by our inability to recover plants with major reductions in the level of DNA methylation (Li *et al*. 2014). Forward genetic screens have uncovered a number of components of the RNA-directed DNA methylation (RdDM) machinery as playing critical roles in maintenance of silenced paramutant states (Dorweiler *et al*. 2000; Jr *et al*. 2009) or transgene silencing (McGinnis *et al*. 2006). These mutants have substantial effects on CHH methylation in maize but have minimal effect on genome-wide levels of CG or CHG methylation (Li *et al*. 2014). Reverse-genetic analyses have identified loss-of-function alleles for a number of other genes predicted to play important roles in DNA methylation but the only single mutants with significant effects on genome-wide DNA methylation are the CMT genes of maize, *Zmet2* and *Zmet5 (Li et al. 2014)*. In this study we have documented how these subtle perturbations of the maize methylome affect the transcriptome in order to find genes subject to epigenetic regulation.

The effects of mutations in RdDM or CMT genes in maize are quite limited. Our evidence suggests that there is little effect on the overall transcriptome of these plants. This might be expected given the limited effect on overall plant phenotype for each of these mutations. A recent study found that *rmr6* (allelic to *mop3*) mutants exhibited transcription changes from a larger portion of the genome but much of this was associated with increased transcriptional ‘noise’ at lowly expressed regions (Forestan *et al*. 2017). However, there are sets of genes with clear changes in expression in each of the mutant lines in our study. Similarly, a directed analysis of differential expression in *rmr6* found a smaller set of genes with significant changes (Forestan *et al*. 2017). While relatively few genes exhibit major changes in expression there are a significant number of genes that exhibit similar expression changes in multiple RdDM or CMT mutants. These findings are compatible with the concept that there are a small number of genes in the maize genome that have epigenetic regulation that is solely dependent upon RdDM or CMT mediated regulation. It is likely that a much larger number of genes are redundantly regulated by the RdDM and CMT pathway along with MET1 mediated CG methylation.

Many of the genes that are up-regulated in CMT mutants exhibit high levels of CHG methylation. The CMT mutants reduce this methylation and allow for increased expression. Previous studies noted that the genes sensitive to *zmet2-m1* mutations varied in different maize inbreds (Makarevitch *et al*. 2007). This prompted us to investigate whether the genes that are up-regulated in CMT mutants might exhibit natural variation for DNA methylation levels. Many of the genes that are up-regulated in CMT mutants have CHG DMRs nearby and many of these exhibit variable levels of expression among maize genotypes that is negatively correlated with CHG methylation levels. This suggests epiallelic diversity for targets of CMT-mediated gene silencing. If these changes in expression lead to phenotypic variation, plant breeders are likely able to select for preferred epigenetic states. However, it would also be possible to introduce novel epigenetic variation through reductions of CHG methylation.

DNA methylation is often considered to play a primary role in maintaining genome integrity by silencing transposable elements. Indeed, there are clear examples of release of transposon silencing in mutants affecting DNA methylation in *Arabidopsis* (Miura *et al*. 2001; Mirouze *et al*. 2009) and maize (Lisch *et al*. 2002; Jia *et al*. 2009). However, generating a complete understanding of transposon expression is complicated by the highly repetitive nature of transposable elements. In order to survey expression using RNAseq most researchers focus on unique mapping reads to ensure that expression is accurately attributed to the proper genomic locus. In this study we elected to primarily focus on TE families rather than individual elements and we utilized an approach that allowed for the combined use of unique and multiple-mapping reads to assess TE family expression. We did find evidence that a number of TE families require RdDM and/or CMT for silencing. There were few distinguishing features about these TEs relative to others making it unclear why the silencing of these families was easily released in these mutants. For several families we were able to document evidence for specific release of silencing of a single member of the family while in other cases we found that multiple members of the same family were reactivated.

This study defines a set of genes and TE families that are regulated by DNA methylation. The silencing of these genes and TEs relies solely upon RdDM or CMT based epigenetic regulation. These loci provide important insights into the mechanisms that allow for epigenetic regulation and the natural variation for epigenetic regulation in maize.

## ACKNOWLEDGEMENTS

We thank Peter Hermanson and Jonathan Giesler for technical assistance and Jonathan Gent for helpful discussions and feedback. Michelle Stitzer and Jeffrey Ross-Ibarra provided helpful discussions and transposon resources. Funding for this work was grants from USDA-NIFA2016-67013-24747 to N.M.S. and the National Science Foundation IOS-1237931 to N.M.S. and M.W.V. Q.L. was supported by The National Key Research and Development Program of China (2016YFD0101003).

## SUPPLEMENTAL MATERIALS

Supplemental Figures S1-S4: attached as separate document

Table S1: Complete list of datasets used in this study.

Table S2: Expression values, differential expression calls, list assignments, and DMR calls for all genes.

Table S3: Expression values, differential expression calls, list assignments, and descriptions for all TE families.

Table S4: Unique mapping read counts and descriptors for each unique transposable element.

## REFERENCES

Alleman M., Sidorenko L., McGinnis K., Seshadri V., Dorweiler J. E., et al., 2006 An RNA-dependent RNA polymerase is required for paramutation in maize. Nature 442: 295–298.

Anders S., Pyl P. T., Huber W., 2015 HTSeq–a Python framework to work with high-throughput sequencing data. Bioinformatics 31: 166–169.

Barber W. T., Zhang W., Win H., Varala K. K., Dorweiler J. E., et al., 2012 Repeat associated small RNAs vary among parents and following hybridization in maize. Proc. Natl. Acad. Sci. U. S. A. 109: 10444–10449.

Bouyer D., Kramdi A., Kassam M., Heese M., Schnittger A., et al., 2017 DNA methylation dynamics during early plant life. Genome Biol. 18: 179.

Brink R. A., 1956 A Genetic Change Associated with the R Locus in Maize Which Is Directed and Potentially Reversible. Genetics 41: 872–889.

Cavrak V. V., Lettner N., Jamge S., Kosarewicz A., Bayer L. M., et al., 2014 How a retrotransposon exploits the plant’s heat stress response for its activation. PLoS Genet. 10: e1004115.

Chandler V. L., Walbot V., 1986 DNA modification of a maize transposable element correlates with loss of activity. Proc. Natl. Acad. Sci. U. S. A. 83: 1767–1771.

Chandler V. L., 2007 Paramutation: from maize to mice. Cell 128: 641–645.

Chomet P. S., Wessler S., Dellaporta S. L., 1987 Inactivation of the maize transposable element Activator (Ac) is associated with its DNA modification. EMBO J. 6: 295–302.

Dong X., Zhang M., Chen J., Peng L., Zhang N., et al., 2017 Dynamic and Antagonistic Allele-Specific Epigenetic Modifications Controlling the Expression of Imprinted Genes in Maize Endosperm. Mol. Plant 10: 442–455.

Dorweiler J. E., Carey C. C., Kubo K. M., Hollick J. B., Kermicle J. L., et al., 2000 Mediator of Paramutation1 is Required for Establishment and Maintenance of Paramutation at Multiple Maize Loci. Plant Cell 12: 2101–2118.

Eichten S. R., Swanson-Wagner R. A., Schnable J. C., Waters A. J., Hermanson P. J., et al., 2011 Heritable epigenetic variation among maize inbreds. PLoS Genet. 7: e1002372.

Eichten S. R., Briskine R., Song J., Li Q., Swanson-Wagner R., et al., 2013 Epigenetic and genetic influences on DNA methylation variation in maize populations. Plant Cell 25: 2783–2797.

Erhard K. F. Jr, Stonaker J. L., Parkinson S. E., Lim J. P., Hale C. J., et al., 2009 RNA polymerase IV functions in paramutation in Zea mays. Science 323: 1201–1205.

Erhard K. F. Jr, Talbot J.-E. R. B., Deans N. C., McClish A. E., Hollick J. B., 2015 Nascent transcription affected by RNA polymerase IV in Zea mays. Genetics 199: 1107–1125.

Forestan C., Aiese Cigliano R., Farinati S., Lunardon A., Sanseverino W., et al., 2016 Stress-induced and epigenetic-mediated maize transcriptome regulation study by means of transcriptome reannotation and differential expression analysis. Sci. Rep. 6: 30446.

Forestan C., Farinati S., Aiese Cigliano R., Lunardon A., Sanseverino W., et al., 2017 Maize RNA PolIV affects the expression of genes with nearby TE insertions and has a genome-wide repressive impact on transcription. BMC Plant Biol. 17: 161.

Gehring M., 2013 Genomic imprinting: insights from plants. Annu. Rev. Genet. 47: 187–208.

Gent J. I., Ellis N. A., Guo L., Harkess A. E., Yao Y., et al., 2013 CHH islands: de novo DNA methylation in near-gene chromatin regulation in maize. Genome Res. 23: 628–637.

Gent J. I., Madzima T. F., Bader R., Kent M. R., Zhang X., et al., 2014 Accessible DNA and relative depletion of H3K9me2 at maize loci undergoing RNA-directed DNA methylation. Plant Cell 26: 4903–4917.

Gouil Q., Baulcombe D. C., 2016 DNA Methylation Signatures of the Plant Chromomethyltransferases. PLoS Genet. 12: e1006526.

Hale C. J., Stonaker J. L., Gross S. M., Hollick J. B., 2007 A novel Snf2 protein maintains trans-generational regulatory states established by paramutation in maize. PLoS Biol. 5: 2156–2165.

Hofmeister B. T., Lee K., Rohr N. A., Hall D. W., Schmitz R. J., 2017 Stable inheritance of DNA methylation allows creation of epigenotype maps and the study of epiallele inheritance patterns in the absence of genetic variation. Genome Biol. 18: 155.

Hollick J. B., Kermicle J. L., Parkinson S. E., 2005 Rmr6 maintains meiotic inheritance of paramutant states in Zea mays. Genetics 171: 725–740.

Hollick J. B., 2017 Paramutation and related phenomena in diverse species. Nature reviews.Genetics 18: 5–23.

Hu L., Li N., Xu C., Zhong S., Lin X., et al., 2014 Mutation of a major CG methylase in rice causes genome-wide hypomethylation, dysregulated genome expression, and seedling lethality. Proc. Natl. Acad. Sci. U. S. A. 111: 10642–10647.

Jia Y., Lisch D. R., Ohtsu K., Scanlon M. J., Nettleton D., et al., 2009 Loss of RNA-dependent RNA polymerase 2 (RDR2) function causes widespread and unexpected changes in the expression of transposons, genes, and 24-nt small RNAs. PLoS Genet. 5: e1000737.

Jiao Y., Peluso P., Shi J., Liang T., Stitzer M. C., et al., 2017 Improved maize reference genome with single-molecule technologies. Nature 546: 524–527.

Jr K. F. E., Stonaker J. L., Parkinson S. E., Lim J. P., Hale C. J., et al., 2009 RNA polymerase IV functions in paramutation in Zea mays. Science 323: 1201–1205.

Kawakatsu T., Stuart T., Valdes M., Breakfield N., Schmitz R. J., et al., 2016 Unique cell-type-specific patterns of DNA methylation in the root meristem. Nature plants 2: 16058.

Kawakatsu T., Nery J. R., Castanon R., Ecker J. R., 2017 Dynamic DNA methylation reconfiguration during seed development and germination. Genome Biol. 18: 171.

Kermicle J. L., 1970 Dependence of the R-Mottled Aleurone Phenotype in Maize on Mode of Sexual Transmission. Genetics 66: 69–85.

Kermicle J. L., Alleman M., 1990 Gametic imprinting in maize in relation to the angiosperm life cycle. Development (Cambridge, England).Supplement: 9–14.

Kim D., Pertea G., Trapnell C., Pimentel H., Kelley R., et al., 2013 TopHat2: accurate alignment of transcriptomes in the presence of insertions, deletions and gene fusions. Genome Biol. 14: R36–2013–14–4–r36.

Li Q., Eichten S. R., Hermanson P. J., Zaunbrecher V. M., Song J., et al., 2014 Genetic perturbation of the maize methylome. Plant Cell 26: 4602–4616.

Li Q., Song J., West P. T., Zynda G., Eichten S. R., et al., 2015a Examining the causes and consequences of context-specific differential DNA methylation in maize. Plant Physiol. 168: 1262–1274.

Li Q., Gent J. I., Zynda G., Song J., Makarevitch I., et al., 2015b RNA-directed DNA methylation enforces boundaries between heterochromatin and euchromatin in the maize genome. Proc. Natl. Acad. Sci. U. S. A. 112: 14728–14733.

Lisch D., Carey C. C., Dorweiler J. E., Chandler V. L., 2002 A mutation that prevents paramutation in maize also reverses Mutator transposon methylation and silencing. Proc. Natl. Acad. Sci. U. S. A. 99: 6130–6135.

Love M. I., Huber W., Anders S., 2014 Moderated estimation of fold change and dispersion for RNA-seq data with DESeq2. Genome Biol. 15: 550.

Madzima T. F., Huang J., McGinnis K. M., 2014 Chromatin structure and gene expression changes associated with loss of MOP1 activity in Zea mays. Epigenetics 9: 1047–1059.

Makarevitch I., Stupar R. M., Iniguez A. L., Haun W. J., Barbazuk W. B., et al., 2007 Natural Variation for Alleles Under Epigenetic Control by the Maize Chromomethylase Zmet2. Genetics 177: 749–760.

Martínez G., Slotkin R. K., 2012 Developmental relaxation of transposable element silencing in plants: functional or byproduct? Curr. Opin. Plant Biol. 15: 496–502.

McGinnis K. M., Springer C., Lin Y., Carey C. C., Chandler V., 2006 Transcriptionally silenced transgenes in maize are activated by three mutations defective in paramutation. Genetics 173: 1637–1647.

Mirouze M., Reinders J., Bucher E., Nishimura T., Schneeberger K., et al., 2009 Selective epigenetic control of retrotransposition in Arabidopsis. Nature 461: 427–430.

Miura A., Yonebayashi S., Watanabe K., Toyama T., Shimada H., et al., 2001 Mobilization of transposons by a mutation abolishing full DNA methylation in Arabidopsis. Nature 411: 212–214.

Narsai R., Gouil Q., Secco D., Srivastava A., Karpievitch Y. V., et al., 2017 Extensive transcriptomic and epigenomic remodelling occurs during Arabidopsis thaliana germination. Genome Biol. 18: 172.

Papa C. M., Springer N. M., Muszynski M. G., Meeley R., Kaeppler S. M., 2001 Maize chromomethylase Zea methyltransferase2 is required for CpNpG methylation. Plant Cell 13: 1919–1928.

Quinlan A. R., Hall I. M., 2010 BEDTools: a flexible suite of utilities for comparing genomic features. Bioinformatics 26: 841–842.

Regulski M., Lu Z., Kendall J., Donoghue M. T., Reinders J., et al., 2013 The maize methylome influences mRNA splice sites and reveals widespread paramutation-like switches guided by small RNA. Genome Res. 23: 1651–1662.

Rowley M. J., Rothi M. H., Böhmdorfer G., Kuciński J., Wierzbicki A. T., 2017 Long-range control of gene expression via RNA-directed DNA methylation. PLoS Genet. 13: e1006749.

Sloan A. E., Sidorenko L., McGinnis K. M., 2014 Diverse gene-silencing mechanisms with distinct requirements for RNA polymerase subunits in Zea mays. Genetics 198: 1031–1042.

Song J., Zynda G., Beck S., Springer N. M., Vaughn M. W., 2016 Bisulfite Sequence Analyses Using CyVerse Discovery Environment: From Mapping to DMRs. In: Current Protocols in Plant Biology, John Wiley & Sons, Inc.

Wang P., Xia H., Zhang Y., Zhao S., Zhao C., et al., 2015 Genome-wide high-resolution mapping of DNA methylation identifies epigenetic variation across embryo and endosperm in Maize (Zea may). BMC Genomics 16: 21.

Wicker T., Sabot F., Hua-Van A., Bennetzen J. L., Capy P., et al., 2007 A unified classification system for eukaryotic transposable elements. Nature reviews.Genetics 8: 973–982.

Williams B. P., Pignatta D., Henikoff S., Gehring M., 2015 Methylation-sensitive expression of a DNA demethylase gene serves as an epigenetic rheostat. PLoS Genet. 11: e1005142.

Woodhouse M. R., Freeling M., Lisch D., 2006 The mop1 (mediator of paramutation1) mutant progressively reactivates one of the two genes encoded by the MuDR transposon in maize. Genetics 172: 579–592.

Xi Y., Li W., 2009 BSMAP: whole genome bisulfite sequence MAPping program. BMC Bioinformatics 10: 232.

Yamauchi T., Johzuka-Hisatomi Y., Terada R., Nakamura I., Iida S., 2014 The MET1b gene encoding a maintenance DNA methyltransferase is indispensable for normal development in rice. Plant Mol. Biol. 85: 219–232.

